# Insights into the molecular mechanisms of cell fate decision making processes from chromosome structural dynamics

**DOI:** 10.1101/2021.05.09.443292

**Authors:** Xiakun Chu, Jin Wang

## Abstract

Cell state transitions or cell fate decision making processes, such as cell development and cell pathological transformation, are believed to be determined by the regulatory network of genes, which intimately depend on the structures of chromosomes in the cell nucleus. The high temporal resolution picture of how chromosome reorganizes its 3D structure during the cell state transitions is the key to understanding the mechanisms of these fundamental cellular processes. However, this picture is still challenging to acquire at present. Here, we studied the chromosome structural dynamics during the cell state transitions among the pluripotent embryonic stem cell (ESC), the terminally differentiated normal cell and the cancer cell using landscape-switching model implemented in the molecular dynamics simulation. We considered up to 6 transitions, including differentiation, reprogramming, cancer formation and reversion. We found that the pathways can merge at certain stages during the transitions for the two processes having the same destination as the ESC or the normal cell. Before reaching the merging point, the two pathways are cell-type-specific. The chromosomes at the merging points show high structural similarity to the ones at the final cell states in terms of the contact maps, TADs and compartments. The post-merging processes correspond to the adaption of the chromosome global shape geometry through the chromosome compaction without significantly disrupting the contact formation. On the other hand, our detailed analysis showed no merging point for the two cancer formation processes initialized from the ESC and the normal cell, implying that cancer progression is a complex process and may be associated with multiple pathways. Our results draw a complete molecular picture of cell development and cancer at the dynamical chromosome structural level, and help our understanding of the molecular mechanisms of cell fate decision making processes.

## 1 Introduction

Cell state transition or cell fate decision making, is one of the fundamental cell events in forming multicellular organisms. The transformation of the normal cell to the cancer cell, which is a typical cell fate transition in pathology, leads to the development of a tumor. The transition processes between distinct cell phenotypic states are believed to be governed by the underlying gene regulatory networks [1, 2]. As the scaffold for the genome function, the chromosome folds into a 3D organization to underpin the gene expressions, which are cell-type-specific. During the transition, the cell undergoes phenotypic switching determined by the complex gene network, which is supported by the large-scale chromosome structural rearrangement in favor of the specific gene expression pattern towards the destined cell state [3]. It has been well recognized that the 3D chromosome structure plays important roles in regulating cell development [4] and cancer [5]. However, our understanding of the abovementioned cell fate decision making processes at the molecular level through chromosome structural dynamics is restricted by the limited experimental approaches, which are often confronted with difficulties in covering the large spatiotemporal scales of the chromosome systems in the cell nucleus.

Currently, the chromosome structural determination is largely performed by the Hi-C techniques and the results are generally presented in terms of an ensemble-averaged contact map [6, 7]. Hi-C experiments measure the frequency of contact formed by all chromosomal loci pairs across the genome, so a high (low) contact frequency formed by two chromosomal loci corresponds to a close (far away) spatial distance and implies a strong (weak) physical interaction between these two loci. Careful analyses on the Hi-C contact map further revealed that the chromosome folds into a hierarchical organization from the multi-megabase compartments [6, 7] to the sub-megabase topologically associating domains (TADs) [8, 9]. These two chromosome structural features have been recognized to have significant impacts on gene expressions [10, 11, 12, 13]. Our knowledge of the chromosome 3D structure and its roles in gene regulation has been significantly increased by the Hi-C techniques. However, Hi-C data only provides the static description of the chromosome organization within one cell state, the dynamical changes of the chromosome structure during the transition between the two cell states are still inaccessible.

Here, we applied the landscape-switching model to study the chromosome structural dynamics during the cell state transitions among the embryonic stem cell (ESC), the normal and cancer cell (See **“Materials and Methods”**). Overall, there are 6 transition processes, including the ESC differentiation to the normal and cancer cell, the normal and cancer cell reprogramming to the ESC, and the cancer cell formation from the normal cell and reversion to the normal cell. The system presents a complete circuitry of cell fate decision making processes, including the normal and pathological developments of cells. The experimental Hi-C contact maps have revealed significant differences in chromosome structures among the ESC, the normal and cancer cell [8, 14]. This feature implies that there are large-scale chromosome structural rearrangements during the cell state transition among these 3 cell states. In our simulations, we constantly monitored the chromosome structures evolving along the time during the cell state transitions and quantified the transition pathways in terms of Hi-C contact map, chromosome structural shape geometry and the spatial distribution of loci on the chromosome.

The quantified pathways of chromosome structural dynamics draw a molecular-level picture of how the cell state transitions or cell fate decision making processes occur. We observed early bifurcation events in two of the cell state transition processes when the initial cell states are the same, such as the ESC differentiation and the processes of deforming the normal and cancer cells. For the two processes with the same destination, the pathways can merge at certain stages, after which the chromosome has to undergo further structural compaction to reach the final state. Detailed structural analyses revealed that the two transition processes for forming cancer cell never overlap, though they can share a similar chromosome structural compaction route. This implies the complex and multiple-route characteristics of cancer formation. Our results suggested that the cell transition processes with the same cell state as the destination for the ESC and normal cell, are made up of an initial cell-type-specific stage followed by a common-route stage. Our study provides a comprehensive understanding of the cell fate decision making processes through chromosome structural dynamics.

## 2 Results

### 2.1 Dynamical chromosome structural rearrangements during differentiation, reprogramming and cancer formation

We calculated the simulated Hi-C contact map evolving in time. We observed that the chromosome dynamically rearranges its structure during the cell state transitions in terms of gradually adapting the Hi-C contact map from the initial cell state towards the final cell state. Interestingly, it appears that the evolutions of the contact maps do not follow the same routes for the forward and reverse transitions between any two cells. This indicates that both the differentiation and reprogramming processes, as well as the cancer formation and reversion processes, are irreversible from the chromosome structural perspective. The irreversibility at the chromosome structural-level can further influence the irreversible gene regulation pathways that determine the non-equilibrium cell state decision-making processes [15, 16].

The differentiation processes starting from the ESC should bifurcate during the transitions towards the normal and the cancer cell, while the normal and the cancer cell reprogramming processes should merge during the transitions towards the ESC. To gain insights into how the bifurcation and merging of the pathways occur from the chromosome structural dynamics, we calculated the correlation coefficient of determination *R*^2^ between the two contact probability matrices in time (Figure 1B and 1C). For reprogramming, it appears that the pathways of the normal and cancer cell reprogramming to the ESC (Repr_N_ and Repr_C_) start to merge quite early at *∼*0.1*τ* with high values of *R*^2^ (Figure 1B, *upper panel*). At *∼* 0.1*τ*, the chromosomes have very different contact formations from either the ESC and final state (Figure 1B, *lower panel*). Interestingly, we observed over-expanded chromosome structures with respect to the one at the ESC formed during the normal and the cancer cell reprogramming processes. Besides, we see that the pathways of reprogramming start to merge before forming the over-expanding chromosome conformation. For differentiation, the two transitions towards the normal and cancer cell (Diff_N_ and Diff_C_) bifurcate at *∼* 2*τ* (Figure 1C, *upper panel*), where the chromosome has different structures from that at the ESC and is also deviated from those at the final cell states (Figure 1C, *lower panel*).

**Figure 1:**
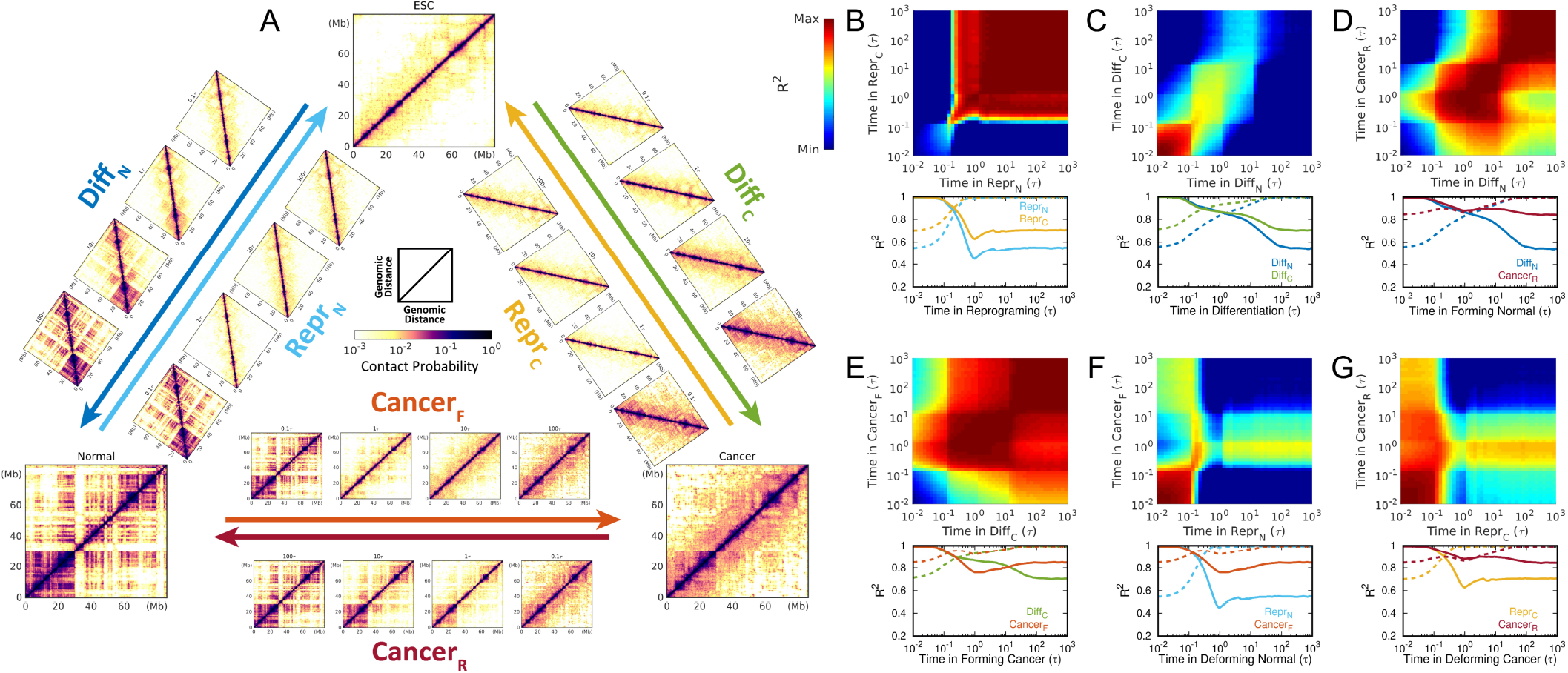
The chromosome structural rearrangements during cell fate decision making among the ESC, the normal and cancer cell. (A) The 6 cell state transition processes associated with chromosome structural dynamics. The chromosome Hi-C contact maps of the ESC, the normal and cancer cell, as well as the cell states at the typical time points (0.1*τ*, 1*τ*, 10*τ*, 100*τ*) during each transition are shown. The correlation maps between the time-course Hi-C contact maps of two transitions during (B) reprogramming from the normal (Repr_N_) and cancer cell (Repr_C_) to the ESC processes, (C) the ESC differentiation to the normal (Diff_N_) and cancer cell (Diff_C_) processes, (D) forming the normal cell from the ESC (differentiation, Diff_N_) and the cancer cell (cancer reversion, Cancer_R_) processes, (E) forming the cancer cell from the ESC (differentiation, Diff_C_) and the cancer cell (cancer formation, Cancer_F_) processes, (F) deforming the normal cell to the ESC (Diff_N_) and the cancer cell (Cancer_F_) processes, (G) deforming the cancer cell to the ESC (Diff_C_) and the normal cell (Cancer_R_) processes. The comparison between process “A” at time *t*^*A*^ = *I* and process “B” at time *t*^*B*^ = *J* is made by calculating the coefficient of determination *R*^2^(*I, J*) between the contact probability *P*_*i j*_ formed by chromosomal loci “i” and 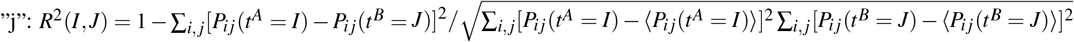. *R*^2^(*I, J*), which measures the similarity of the two Hi-C contact map at time *t*^*A*^ = *I* of process A and *t*^*B*^ = *J* of process B (*R*^2^(*I, J*)=1 corresponds to the identical Hi-C contact maps with *P*_*i j*_ (*t*^*A*^ = *I*)=*P*_*i j*_ (*t*^*B*^ = *J*) and the deviation of *R*^2^(*I, J*) from 1 indicates the degree of difference between these two contact maps.), is shown by 2D plots colored from blue (minimum) to red (maximum) at the upper panels in (A-G). At the lower panels in (A-G), the cell states “J” are selected to be either the initial states (solid lines) or the end states (dashed lines).

Since the destinations of differentiation Diff_N_ and cancer reversion to the normal cell (Cancer_R_) are the same, it implies that these two pathways should merge at certain stages of these two transitions. From Figure 1D, we see that the chromosomes during Diff_N_ and Cancer_R_ transitions become structurally similar at *∼* 1*τ*, where the chromosome has accomplished significant structural rearrangements for forming the normal cell (Figure 1D, *lower panel*). However, a considerable number of the chromosome contact changes after 1*τ* is still seen during Diff_N_ to make the chromosome gradually deviate from the ESC. A similar observation can be found in forming the cancer cell by differentiation Diff_C_ and cancer formation process (Cancer_F_) (Figure 1E). The contact maps of the two pathways become similar at *∼* 1*τ*, where the chromosome with different structures from that at the ESC still has to undergo a certain degree of structural arrangements in order to form the structure at the cancer cell.

In addition to the ESC differentiation, we also studied the bifurcation of the processes with the initial cell states being the normal cell (Repr_N_ and Cancer_F_, Figure 1F) and cancer cell (Repr_C_ and Cancer_R_, Figure 1G), respectively. The two pathways in both of these processes exhibit bifurcation at the early stages of the transitions around 0.2*τ*, which is later than the bifurcation time observed at the ESC differentiation (*∼* 2*τ*). It is worth noting that during Cancer_F_ and Cancer_R_, the chromosome at 0.1-1*τ* shares a certain degree of structural similarity with that after 1*τ* during the reprogramming towards the ESC. This feature has led to an implication that the cell during the transitions between the normal and cancer states increases the degree of the “stemness” from the chromosome structural perspective [17].

### 2.2 Chromosome structural shape changes during differentiation, reprogramming and cancer formation

Using molecular dynamics simulations, we can have the 3D chromosome structural information in addition to the 2D contact map. Here, we introduced three reaction coordinates to describe the geometry of the chromosome structural shape. The radius of gyration *R*_*g*_ is used to describe the chromosome compaction and the aspheric shape parameter Δ is used to measure the extent of anisotropy with deviation from 0 as an indication of deviation from a perfect sphere [18]. The lengths of chromosome structure along the longest principal axis (PA1) and shortest principal axis (PA3) are used to describe the global geometric shape of the chromosome structure.

We observed the over-expanded chromosome structures with respect to the one at the ESC during both the normal and cancer cells reprogramming to the ESC processes (Figure 2A). During both of the reprogramming processes, the chromosome expanding stages are isotropic and reach the over-expanded phase with a similar shape at similar time points (1*τ*). Then, the chromosome undergoes slight compaction to form the chromosome shape at the ESC through the merging pathways with negligible contact probability adaption (Figure 1B). From the chromosome structural perspective, these two reprogramming processes use different routes to form the same over-expanded chromosome structures, implying that the reprogramming pathways are cell-type-specific at the early stage but conserved for different cell types at the late stage.

**Figure 2:**
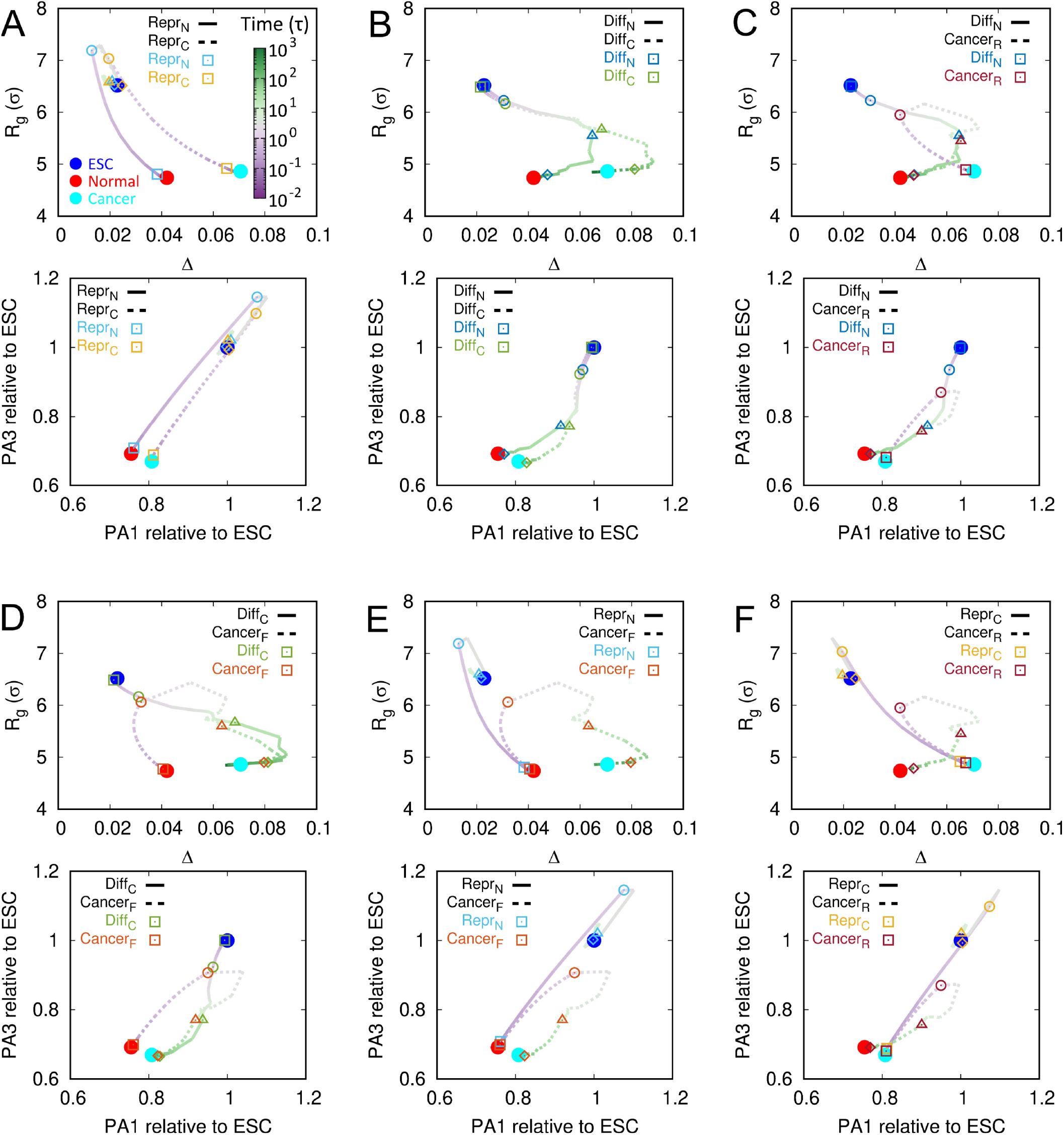
Quantified averaged pathways for chromosome structural shape changes during cell fate decision making among the ESC, the normal and cancer cell. The pathways are shown for (A) differentiation, (B) reprogramming, (C) forming the normal cell, (D) forming the cancer cell, (E) deforming the normal cell and (F) deforming the cancer cell. The reaction coordinates are the radius of gyration (*R*_*g*_) and aspheric shape parameter (Δ) at the upper panels in (A-E), and the length of structural geometric extensions at the longest and shortest principal axes (PA1 and PA3) at the lower panels in (A-E). In each panel, the typical time points in the trajectories are additionally shown as squares (□, t=0.1*τ*), circles (◯, t=1*τ*), triangles (△, t=10*τ*) and diamonds (◊, t=100*τ*). Time is in the logarithmic scale.

For cell differentiation, the chromosome structural pathways with shape changes share the same route towards the normal and cancer cells till *∼* 10*τ* (Figure 2B), when a large degree of contact maps in the destined cell states are established (Figure 1C). From the chromosome contact formation perspective, we observed that the bifurcations of the two differentiation processes start at an earlier time of 1*τ* than that based on the chromosome structural shape-changing pathways (Figure 1C). These features indicate that the geometric chromosome compaction during the ESC differentiation to the normal and cancer cell share the same route for a long duration at the first stage of differentiation, even when the chromosome contact maps within these two transitions have deviated from each other already.

From the chromosome structural shape-changing pathways, we can see that the two processes of forming the normal cell by the ESC differentiation and cancer reversion merge at *∼* 10*τ* (Figure 2C), where the chromosome has a more open structure than that at both the normal and cancer cell. Thus, the merging pathways correspond to chromosome compaction processes associated with forming the chromosome contact maps at the normal cell. It appears that the chromosome contact maps become similar at the very early stages of Diff_N_ and Cancer_R_ around 1*τ* (Figure 1D), so the chromosomes within these two processes can form similar structural shapes. These findings imply that the chromosome contact changes during 1-10*τ* contribute to the increase of the similarity of the chromosome structural shapes during these two processes. A similar observation can be observed for forming the cancer cell by Diff_C_ and Cancer_*F*_. Overall, we found that the transitions between the normal and cancer cells are made up of an initial chromosome structure expansion followed by compaction. The late stages share similar routes with the chromosome geometric compaction pathways during the ESC differentiation.

For the cell state transitions with the same initial states at the normal and cancer cell, the pathways projected onto the chromosome shape parameters bifurcate at the very beginning around 0.1*τ*. In order words, although the chromosome has to undergo an initial structural expansion during the transitions between the normal and cancer cells, the pathways deviate from the ones during reprogramming. The early bifurcation in the chromosome shape-changing pathways is coincident with what was observed from the contact probability perspective (Figure 1F and 1G). This indicates that the arrangements of the contact maps at the early stage of deforming the normal and cancer cell contribute significantly to the specific adaptions of the chromosome structural shape towards the destined cells.

### 2.3 Spatial rearrangement of chromosomal loci during differentiation, reprogramming and cancer formation

The spatial distribution of the loci in chromosome has significant impacts on the transcriptional activity [19, 20]. We calculated the changes of the radial density of the chromosome during the 6 transitions (*ρ*(*r, t*)) and further classified them into the descriptions of the whole loci, loci in compartment A and compartment B (Figure S3-S5). Then we collected *ρ*(*r, t*) for all the processes and performed the principal component analysis (PCA) on all sets of *ρ*(*r, t*). We projected the chromosome structural dynamics during differentiation, reprogramming and cancer onto the first and second principal components (PC1 and PC2) and obtained the trajectories in terms of dynamically arranging the spatial distribution of the chromosome loci (Figure 3).

**Figure 3:**
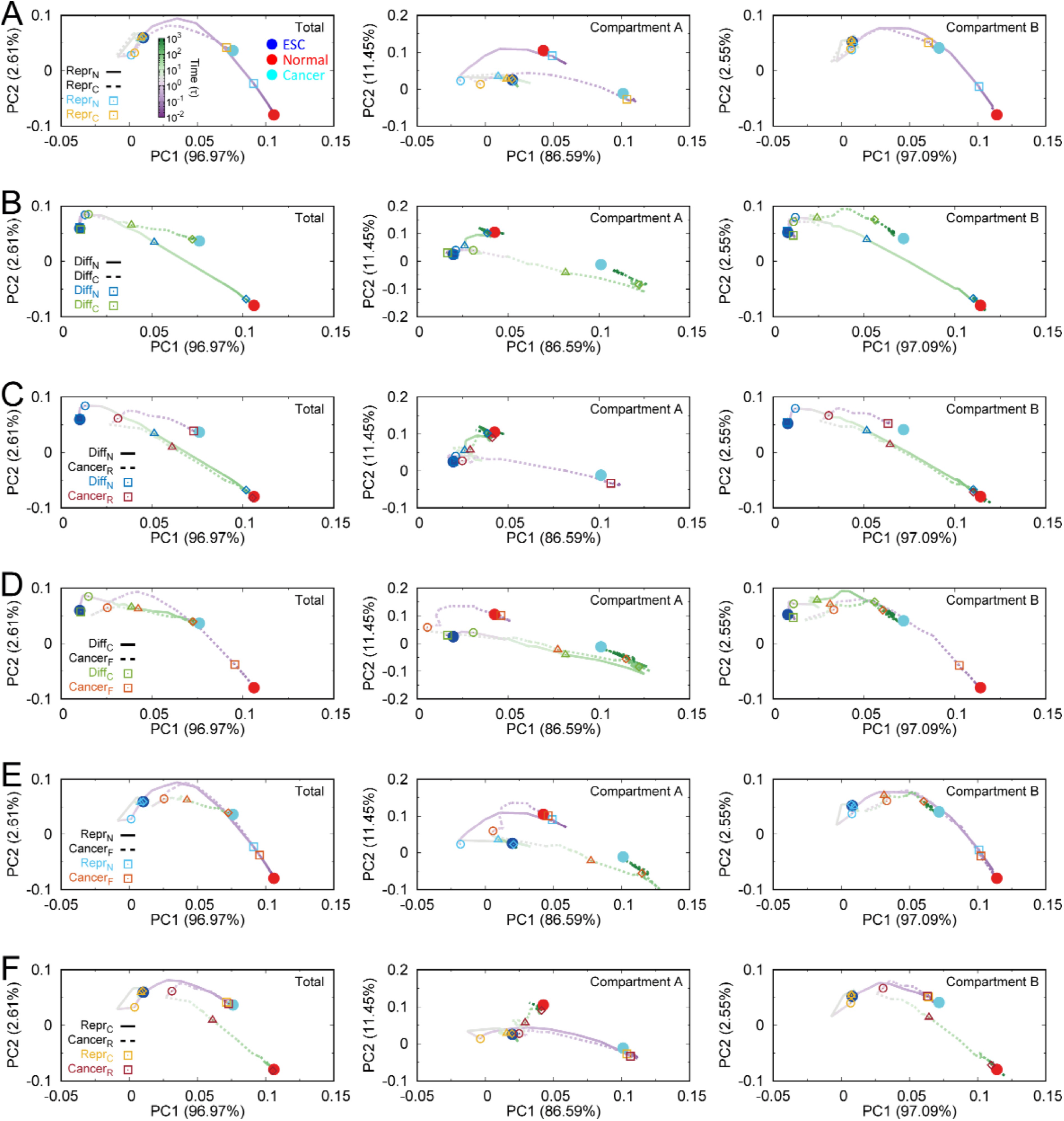
Quantified averaged pathways for the spatial distribution of chromosomal loci changing during cell fate decision making among the ESC, the normal and cancer cell. The pathways are shown for (A) differentiation, (B) reprogramming, (C) forming the normal cell, (D) forming the cancer cell, (E) deforming the normal cell and (F) deforming the cancer cell. The trajectories are projected on the the first and second principal components (PC1 and PC2) of the spatial distribution of chromosomal loci changes for the whole loci (left panel), loci in compartment A (center panel) and loci in compartment B (right panel). In each panel, typical time points in the trajectories are additionally shown as squares (□, t=0.1*τ*), circles (◯, t=1*τ*), triangles (△, t=10*τ*) and diamonds (◊, t=100*τ*). Time is in the logarithmic scale.

Overall, we see that the rearrangements of the spatial distribution of the whole loci share similar trends of the loci in compartment B for all the transitions. During the normal cell reprogramming process, the changes of *ρ*(*r*) of the whole loci and loci in compartment B appear to go through that at the cancer cell state (Figure 3A), while the *ρ*(*r*) of the loci in compartment A can only have overlap with that during cancer cell reprogramming starting at the over-expanding stages. This implies that the distinct reprogramming routes from different cells at the early stage strongly depend on the spatial arrangement of the loci in compartment A. For the ESC differentiation, we see that the bifurcations of the two pathways to the normal and cancer cells start early for both *ρ*(*r*) of the loci in compartment A and compartment B (Figure 3B). This indicates a unique evolution route to the destined terminally differentiated cell for the ESC differentiation.

For forming the normal cell, the pathways of *ρ*(*r*) for differentiation and cancer reversion processes merge (Figure 3C). For the whole loci and loci in compartment B, the merging appears when there is still a long way to form the final *ρ*(*r*) at the normal cell. Interestingly, we found that the merging point for*ρ*(*r*) of loci in compartment A is located near to the position of *ρ*(*r*) at the ESC. Similar trends can be observed in forming the cancer cell from the ESC and the normal cell (Figure 3D). However, there is a slight discrepancy between the *ρ*(*r*) pathways of loci in compartment B for Diff_C_ and Cancer_F_, implying that these two processes for forming the cancer cell may not use the same routine to the spatial repositioning of the chromosomal loci.

We also observed the bifurcation of the pathways from the *ρ*(*r*) between Repr_N_ and Cancer_F_ (Figure 3E), and between Repr_C_ and Cancer_R_ (Figure 3F), similar to that from the chromosome structural shape perspective. However, the elements leading to the bifurcation of the two pathways in deforming the normal cell and the two pathways in deforming the cancer cell are different. For Repr_N_ and Cancer_F_, we see an overlap between these two pathways at the very beginning with *ρ*(*r*) of the whole loci and loci in compartment B. Thus, the evolution of *ρ*(*r*) of the loci in compartment A contributes to the differences of these two pathways. In contrast, for Repr_C_ and Cancer_R_, *ρ*(*r*) of the loci in compartment A shares the same route towards forming the *ρ*(*r*) at the ESC, so the bifurcation of these two pathways is led by the changing *ρ*(*r*) of the loci in compartment B.

### 2.4 Chromosome structures in transient states during differentiation, reprogramming and cancer formation

Based on the above pathways analysis, we can see that the transitions among the differentiation, reprogramming and cancer can merge at certain points. Identifying these points can provide a full picture of how these processes occur from the chromosome structural perspective. To determine these merging points, we first chose a series of time points located on the pathways projected on the chromosome structural shape parameters and PCs of *ρ*, when the two pathways (Repr_N_ and Repr_C_, Diff_N_ and Cancer_R_, Diff_C_ and Cancer_F_) start to merge. Then we calculated the *R*^2^ of the contact maps between the two time points on these two pathways and selected the highest *R*^2^ as the time when the pathways emerge (Figure S6). There are 6 transient states determined as the merging points for reprogramming to the ESC (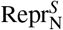 and 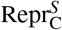), forming the normal cell (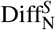 and 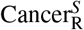) and forming the cancer cell (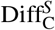 and 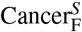). In Figure 4, we can see one transient state is overlapped with the other for determining the merging point. However, it is worth noting that 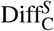 and 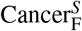 have an apparent separation on pathways of *ρ*(*r*) of loci in compartment B. It implies that the chromosomes at 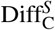 and 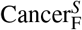 should be structurally different.

**Figure 4:**
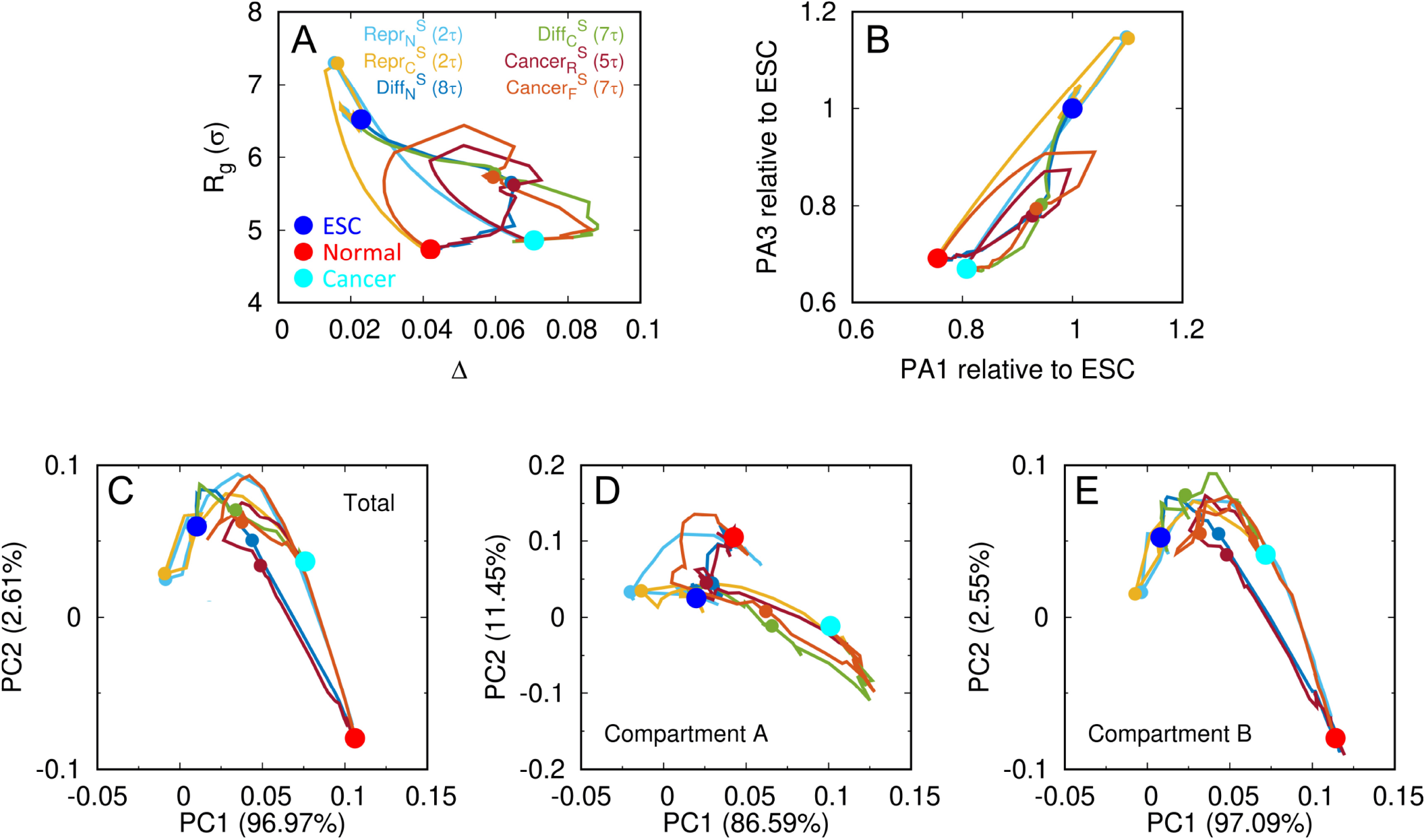
All the averaged trajectories of the 6 processes for differentiation, reprogramming and cancer involved with the transient states projected onto (A) *R*_*g*_ and Δ, (B) extension lengths at PA1 and PA3, and PC1 and PC2 of spatial distribution of chromosome loci for (C) the whole loci, (D) loci in compartment A and (E) loci in compartment B.

The direct comparison of the contact maps between the two transient states at the merging points indicates high similarities of chromosome structural formation with *R*^2^ all equal to 0.99 (Figure 5(A-F)). To examine the similarity of the chromosome structures in terms of the TADs and compartments, we further calculated the signals of the insulation score and compartment profile. We found that the formations of TAD boundaries of the chromosome at the two transient states for one merging point are very similar and close to the ones at the destined cell states (Figure 5G and Figure S8). It is worth noting that chromosomes at the merging point for forming normal cell (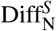 and 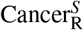) and for forming cancer cell (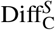 and 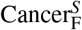) have similar structural shape, but the signals of the TAD boundaries are not strongly correlated. During our simulations, we did not detect significant changes in TAD boundaries (Figure S9), resonating with the experimental evidence that TAD features are mostly conserved in mammals across different cell types and species [8, 21]. For compartments, we found strong correlations of the signals between transient states at the merging point of reprogramming (Repr_N_ and Repr_C_) and forming normal cell (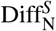 and 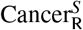), while the compartment profiles are quite different at 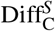 and 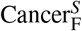 for the merging point in the cancer formation process (Figure 5H and Figure S10). Thus, we can state that the chromosomes at 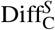 and 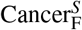 are structurally different, leading to the different compartment formation. The pathways from the ESC and normal cell in forming the cancer cell are not merged based on the chromosome structural formation at the compartment level.

**Figure 5:**
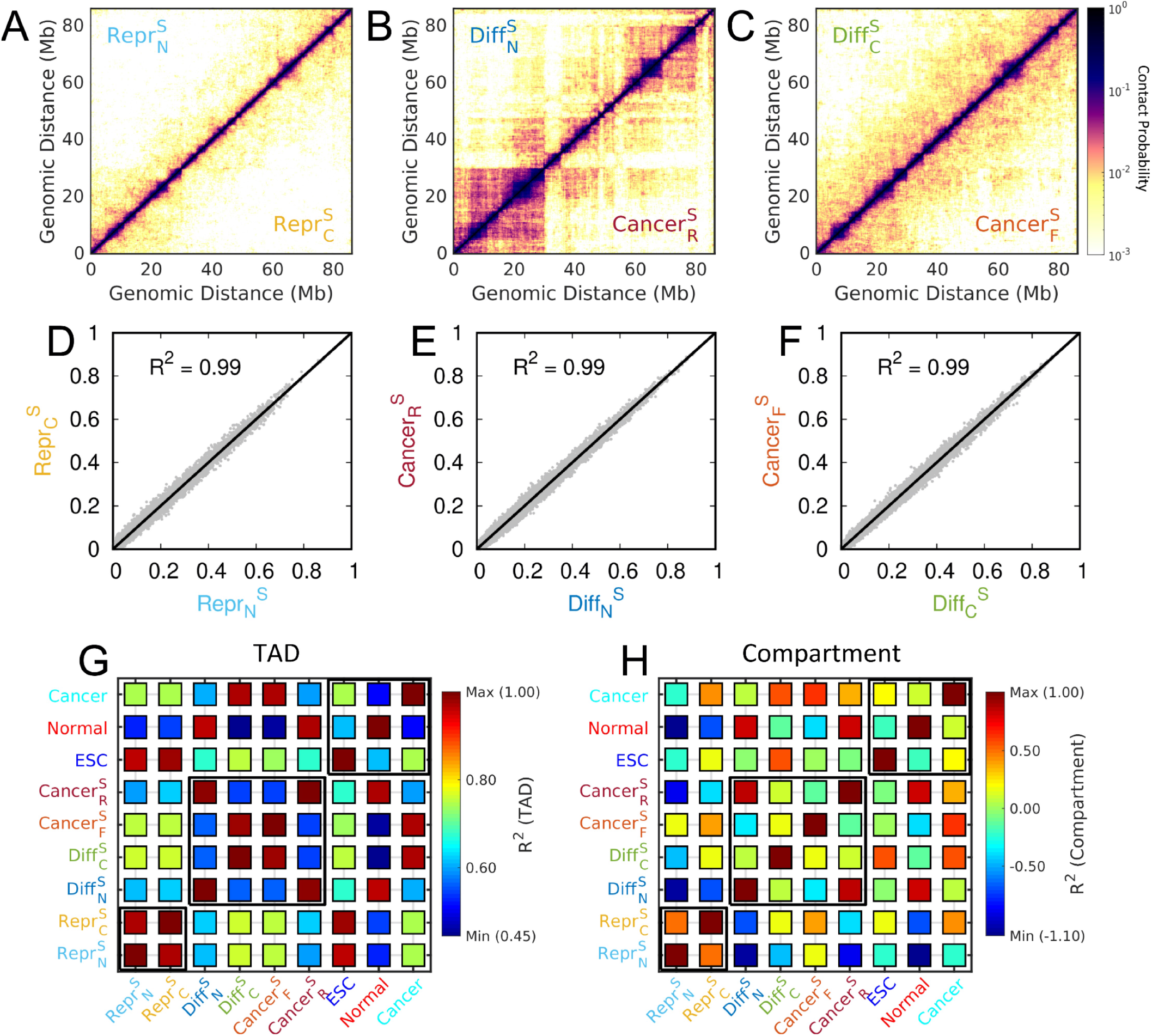
Correlations of chromosome structures between the transient states during cell fate decision making among the ESC, the normal and cancer cell. The contact probability maps for the transient states formed during (A) reprogramming, (B) forming the normal cell and (C) forming the cancer cell. The coefficient of determination *R*^2^s are shown under the corresponding Hi-C contact maps in (D-F). Chromosome structural similarities among the transient states, ESC, normal and cancer cells in terms of (G) TAD and (H) compartment. For TAD, *R*^2^ is calculated based on the insulation score. For compartment, *R*^2^ is calculated based on the compartment profile. The calculation procedures of TAD insulation score and compartment profile can be found in “**Materials and Methods**”.

## 3 Discussion and conclusions

In this work, we studied the chromosome structural dynamics during the transitions between the two different cell states in cell fate decision making among the ESC, the normal and cancer cell. At the chromosomal structural-level, we showed that the forward and reverse cell state transitions between two cell states, such as the differentiation and reprogramming, the cancer formation and reversion, use different routes, leading to irreversible processes. More importantly, we found that the chromosome structural dynamical pathways for the transition towards the same cell can merge prior to the destination. The chromosomes at the merging points are structurally different from those at the corresponding final cell state. Our findings indicate that further chromosome structural arrangements, which are necessary for accomplishing the transition, undergo the same pathway for different processes after passing the merging point.

Based on the chromosome structural dynamical pathways from our analyses, we can draw a full picture of the cell state transitions among the ESC, the normal and cancer cell with more molecular-level details than the one that only contains 6 direct transitions between two cell states at the gene expression level (Figure 6). For reprogramming, the normal and cancer cells are initially converted to the merging point (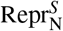 and 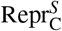), followed by the further transition to the ESC using the same pathway. At the merging point, the chromosomes have a high degree of similarity of the contact probability map with that at the ESC, and slight over-expanded structures compared to the one at the ESC. In other words, the normal and cancer cells have to undergo a long journey of cell-type-specific reprogramming before reaching the merging point, where most of the chromosome interactions at the ESC have been established with an open chromosome structure. We note that the experimental identifications on the transcriptional dynamics trajectories of different somatic cells reprogramming to the iPSC are strongly dependent on the cell types [22]. This appears to be consistent with the first stages of reprogramming processes to the merging point observed in our simulations. A recent integrative transcriptional and epigenomic analysis on the human reprogramming process with a high temporal resolution unprecedentedly uncovered that the gene network established at the late reprogramming process has a signature of pre-implantation stages [23]. During pre-implantation stages, the chromosome undergoes the structural reorganization in establishing the transition from the totipotency to pluripotency state [24]. The nuclear volumes in many mammalian embryos have been found to gradually decrease prior to the blastocyst stages during early embryonic development, leading to a universal chromosome structural compaction process during the pre-implantation stages [25, 26]. Our theoretical predictions on the late reprogramming stage after passing the merging point at the 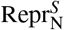 and 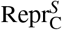 with the over-expanded chromosome conformation are reminiscent of the pre-implantation stages of the reprogramming process [23]. We also found that the cell states, which originate from the normal and cancer cell at the merging point, and the following pathways towards the ESC are conserved regardless of the cell types. This echoes the conserved pre-implantation stages, where the cell has not started the cell-specific differentiation.

**Figure 6:**
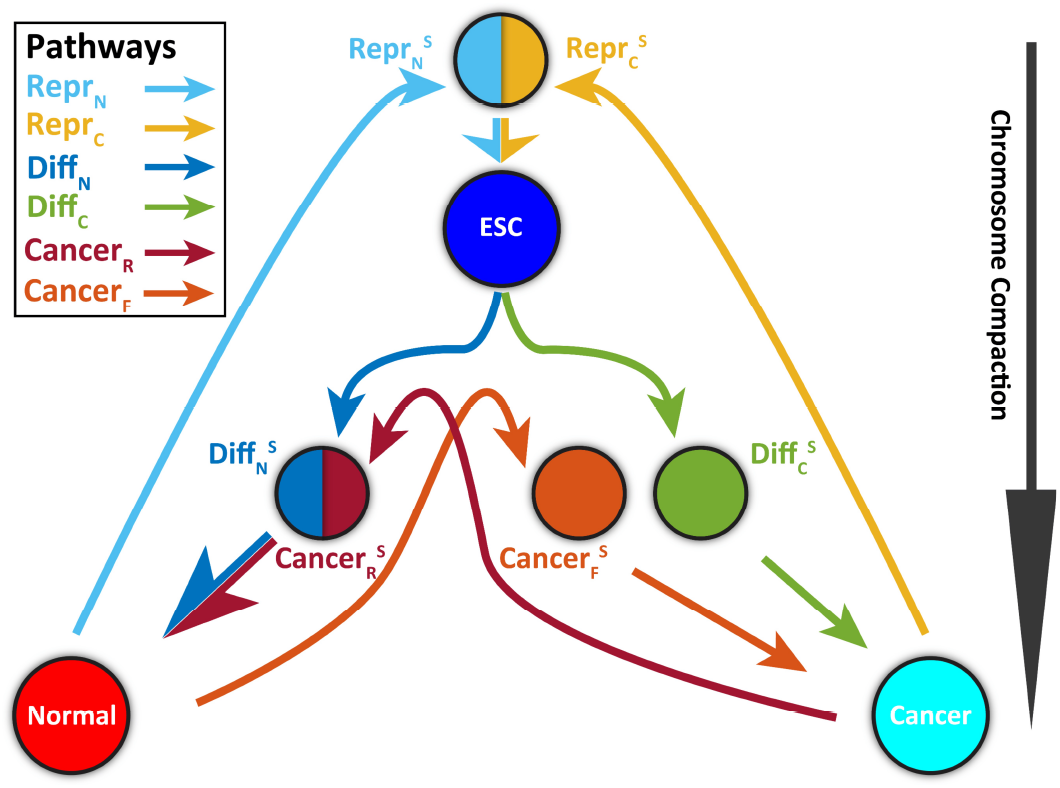
Scheme illustrating the 6 cell state transition processes among the ESC, the normal and cancer cell involved with the transient states from the chromosome structural dynamics perspective. The vertical arrow from the top to the bottom indicates the degree of chromosome compaction.

We also found that the chromosome structural dynamics pathways of the differentiation and cancer reversion processes towards the normal cell are conserved at the late stages of cell state transitions. The chromosomes during these two transitions at the merging point of the pathways (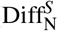 and 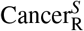) show high structural similarity. In addition, the TAD boundaries and compartments in chromosomes at the 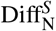 and 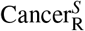 are similar to those at the normal cell. Further compaction of the chromosome from the cell state at the merging point towards the normal cell requires the spatial repositioning of the chromosome loci, while the local chromosome structures and compartment status of the chromosomal loci remain largely unaltered. As the TAD structures and compartment segregations in the chromosome contribute significantly to the gene expressions that determine the cell fate [13, 27, 28], we speculate that the chromosomes at the merging point of the differentiation and cancer reversion transition have accomplished most of the structural arrangements for the functional purposes at the final cell state. In other words, the pathways for forming the normal cell merge at the very late stages of the cell state transition processes.

Interestingly, the two processes of forming the cancer cell from the ESC and normal cell do not share the chromosome structural dynamics pathways, in contrast with either the reprogramming processes or the processes of forming the normal cell. Despite showing similar degrees of global chromosome compaction and the overall chromosome contact map patterns, the chromosomes at the 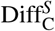 and 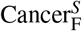 with similar TAD structures exhibit significant differences at the compartment formation. Further changing the compartment status with spatial repositioning of the chromosome loci from 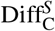 and 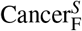 towards the cancer cell is indispensable and undergoes different pathways for the differentiation and cancer formation processes. Therefore, our results show multiple routes for forming the cancer cell from the chromosome structural perspective. The results here focused on the chromosome structural dynamics at the molecular level resonate with our recent theoretical work using the gene regulatory network, which revealed the global mechanisms of cancer with multiple states and paths, underlining the complexity of cancer at the gene level [29].

Our theoretical predictions can be assessed by future biophysical experiments. In this respect, the recently developed time-course single-cell Hi-C experiments can be implemented to measure the chromosome structural rearrangements for the differentiation, reprogramming and cancer processes [30, 31]. The additional Hi-C data on the intermediate states in the processes can also be incorporated into the landscape-switching model, leading to stepwise switching transitions. It is expected that the multistep landscape-switching model can produce more realistic results as more experimental information is incorporated. In summary, the picture drawn here provides a comprehensive and physical understanding of the cell state transition processes among the pluripotent cell, the terminally differentiated cell and the cancer cell at the chromosomal, molecular, and structural levels.

## 4 Materials and Methods

### 4.1 Hi-C data

The Hi-C data of the ESC and the normal cell (IMR90, human fetal lung cell) were downloaded from the Gene Expression Omnibus (GEO) repository archives with accession number GSE35156 [8]. The Hi-C data of the cancer cell (A549, human lung carcinoma cell) was obtained from the ENCODE project with GEO accession number GSE105600 [14]. All the replicas in each dataset of Hi-C data were used and proceeded to the Hi-C Pro software following the standard pipeline [32]. All the Hi-C contact maps were built at 100kb resolution after iterative correction and eigenvector decomposition (ICE) normalization. Here, we focused on the long arm of chromosome 14 (20.5-101.Mb).

### 4.2 Polymer model

The polymer simulation is based on a “beads-on-a-string” model. Each bead, representing a 100kb chromosome segment, is connected by pseudo bonds to the neighboring beads. Thus, the system has a total of 857 beads. We used the finitely extensible nonlinear elastic (FENE) potential to describe the bond stretching [33]. The additional repulsive potential was added between the neighboring beads to avoid the spatial overlap [34]. The angle potential for the consecutive three beads and long-range soft-core potential for non-bonded beads were implemented as suggested previously [35]. Spherical confinement was added to mimic the volume fraction of the chromosome inside the cell nucleus at 10% [36]. The bonded and non-bonded potentials together lead to a typical homopolymer potential and finally generates an equilibrium globule chromosome ensemble, which has no TAD and compartment formation [37]. Details of the polymer model can be found in our previous work [38].

### 4.3 Maximum entropy principle simulation

Hi-C data was additionally incorporated into the polymer potential as the experimental restraint through the maximum entropy principle [39]. The maximum entropy principle generates a chromosome ensemble that can reproduce the experimental Hi-C data with a minimal bias by maximizing the entropy of the system. We performed the maximum entropy principle simulations for the ESC, normal and cancer cell, respectively. The simulations lead to three data-driven potentials, under which the chromosomes form three heterogeneous structural ensembles, which exhibit the experimentally consistent Hi-C contact maps for the ESC, normal and cancer cell, respectively [37, 17]. The potential was further surveyed to be capable of generating the chromosome dynamics that is in good agreement with multiple experiments in many aspects [40, 41], including anomalous diffusion, viscoelasticity, and spatially coherent dynamics [42, 43]. These features suggest that the potential can be regarded as an effective equilibrium landscape that governs both the chromosome structure and dynamics in one cell state.

### 4.4 Landscape-switching model

To explore the chromosome dynamics during the cell state transitions, we used the landscape-switching model, which was developed in our previous work [38, 37, 17]. To connect the landscapes of two cells among the ESC, normal and cancer cell, we performed two sets of molecular dynamics simulations of chromosomes under the potentials of two distinct cell states, separately. After a long duration of simulations when the systems become steady, we switched these two potentials (landscapes), which occurs abruptly to recapitulate the bistable switch between the two gene expression states of two cell states in cell development [44, 45, 46]. From a physical perspective, this implementation resembles an instantaneous energy excitation that drives the system out-of-equilibrium. Then, the chromosomes relaxed on the post-switching potential landscapes and the relaxation processes were referred to as the chromosome dynamics during the transitions between these two cell states. The landscape-switching model has led to non-equilibrium non-adiabatic dynamics, thus it can be used to capture the essence of cell state transitions for the interlandscape crossing events (transition between the steady stable cell states) [47, 48]. In this study, the landscape-switching model leads to 6 transitions indicated in Figure 1A, including the ESC differentiation to the normal cell and cancer cell, the reprogramming from the normal cell and cancer cell, the cancer formation from the normal cell and the cancer reversion to the normal cell. The trajectories of the relaxation processes in the landscape-switching model were extracted and analyzed for describing the chromosome structural dynamics during the cell state transition processes. Details of maximum entropy principle simulation and landscape-switching model can be found in our previous work [38, 37, 17].

### 4.5 Identification of TADs and compartments

The TAD structures were described by the insulation score, which was previously used to determine the boundaries of the TADs [49]. The sliding space (5 *×* 5 beads) was used to calculate the insulation score from the contact map at each cell state. The compartment profiles were calculated by contact probability *P*_obs_/*P*_exp_, which is the ratio between the observed contact probability *P*_obs_ and expected contact probability *P*_exp_ at the 1Mb resolution. Then the principal component analysis (PCA) was performed on the enhanced contact probability map after the ICE normalization. Finally, the first principle component (PC1) was used to represent the compartment profile. The compartment status of the normal cell was determined by the direction of the PC1 values, which was assigned with the gene density (positive to gene-rich and compartment A, negative to gene-poor and compartment B). The direction of the PC1 values of other cell states was determined by calculating the correlation coefficient to the compartment profile (PC1 with the correct direction) of the normal cell.

## Acknowledgement

We acknowledge the support from the National Science Foundation PHY-76066. The authors would like to thank Stony Brook Research Computing and Cyberinfrastructure, and the Institute for Advanced Computational Science at Stony Brook University for access to the high-performance SeaWulf computing system, which was made possible by a $1.4M National Science Foundation grant (#1531492).

