## Supporting Information for "Insights into the molecular mechanisms of cell fate decision making processes from chromosome structural dynamics"

for

### Figures

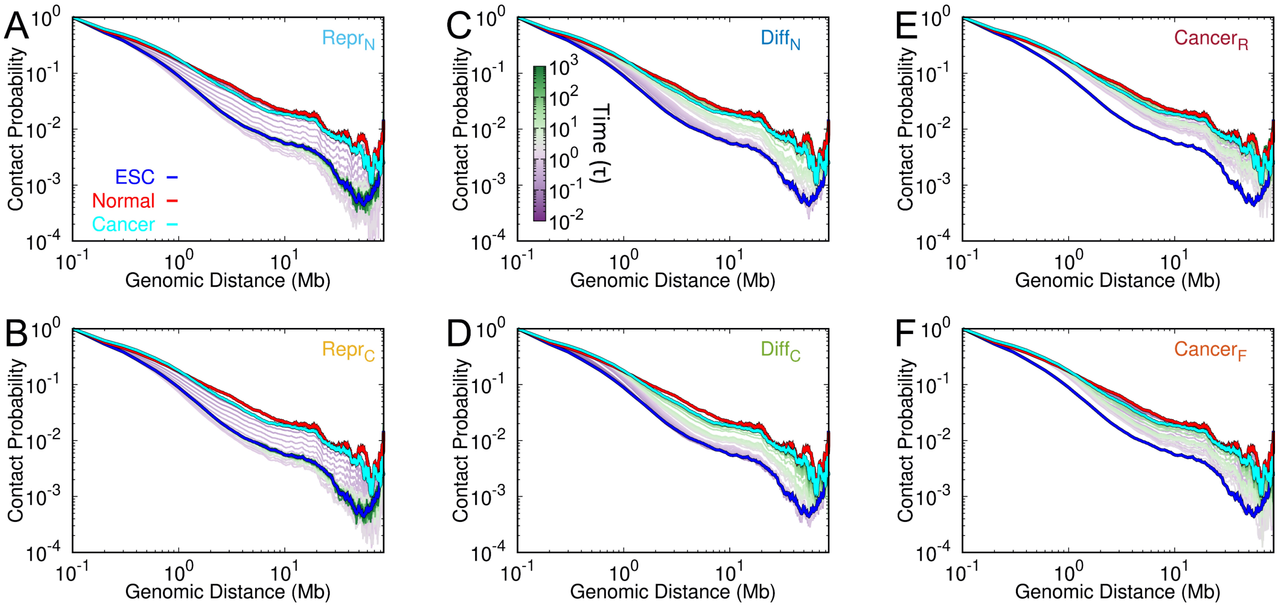


Figure S1: The contact probability $P(l)$ along the genomic distance $l$ during the processes of (A) normal cell reprogramming to ESC, (B) cancer cell reprogramming to ESC, (C) ESC differentiation to normal cell, (D) ESC differentiation to cancer cell, (E) cancer reversion to normal cell and (F) cancer formation from normal cell. Time is in the logarithmic scale. In each panel, the curves $P(l)$ at ESC, normal and cancer are also plotted.

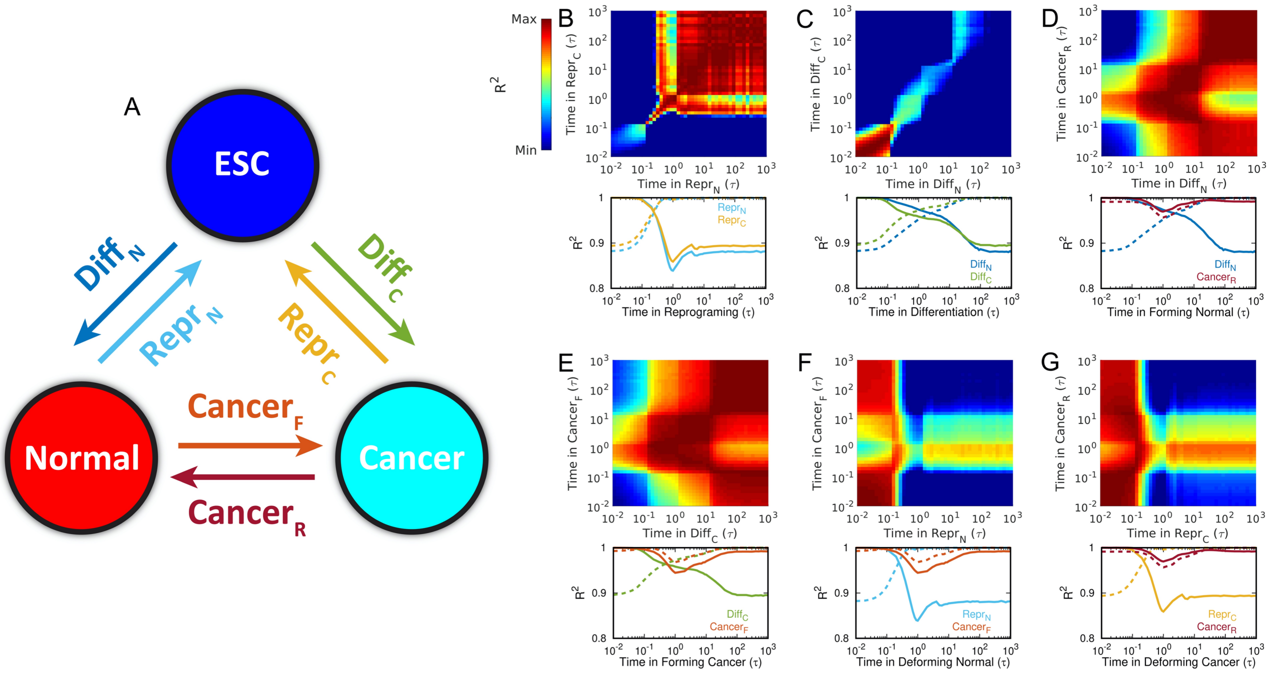


Figure S2: Similar with Figure 1 but for comparing $P(l)$ among the processes of differentiation, reprogramming and cancer.

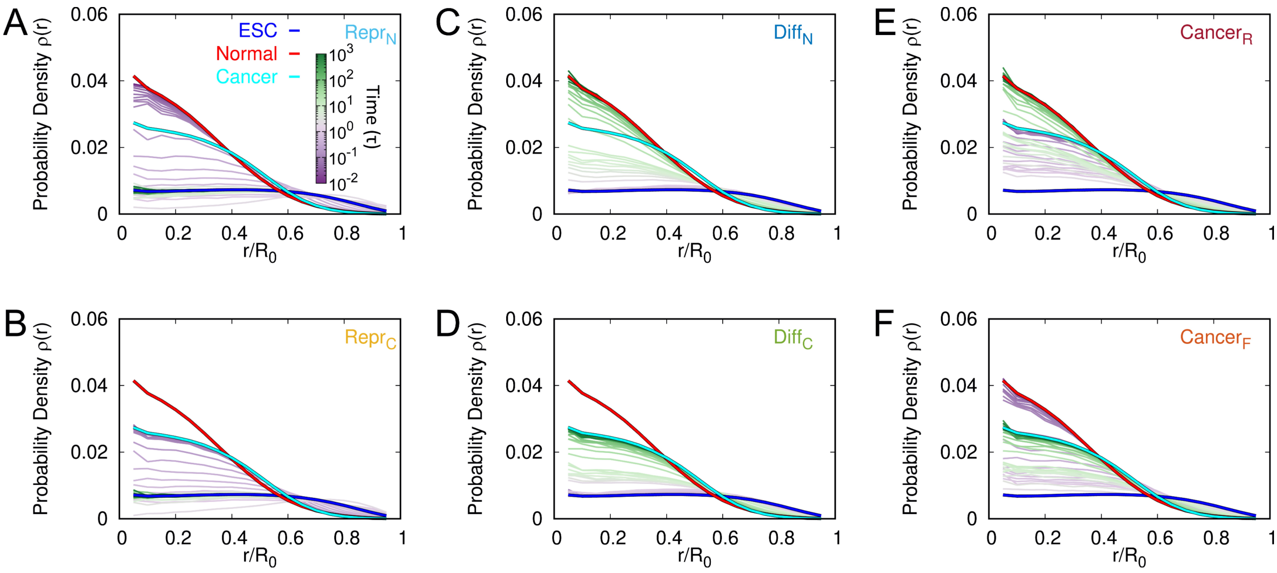


Figure S3: The change of the radial density $\rho(r)$ in the chromosome for the **whole loci** during the processes of (A) normal cell reprogramming to ESC, (B) cancer cell reprogramming to ESC, (C) ESC differentiation to normal cell, (D) ESC differentiation to cancer cell, (E) cancer reversion to normal cell and (F) cancer formation from normal cell. Time is in the logarithmic scale.

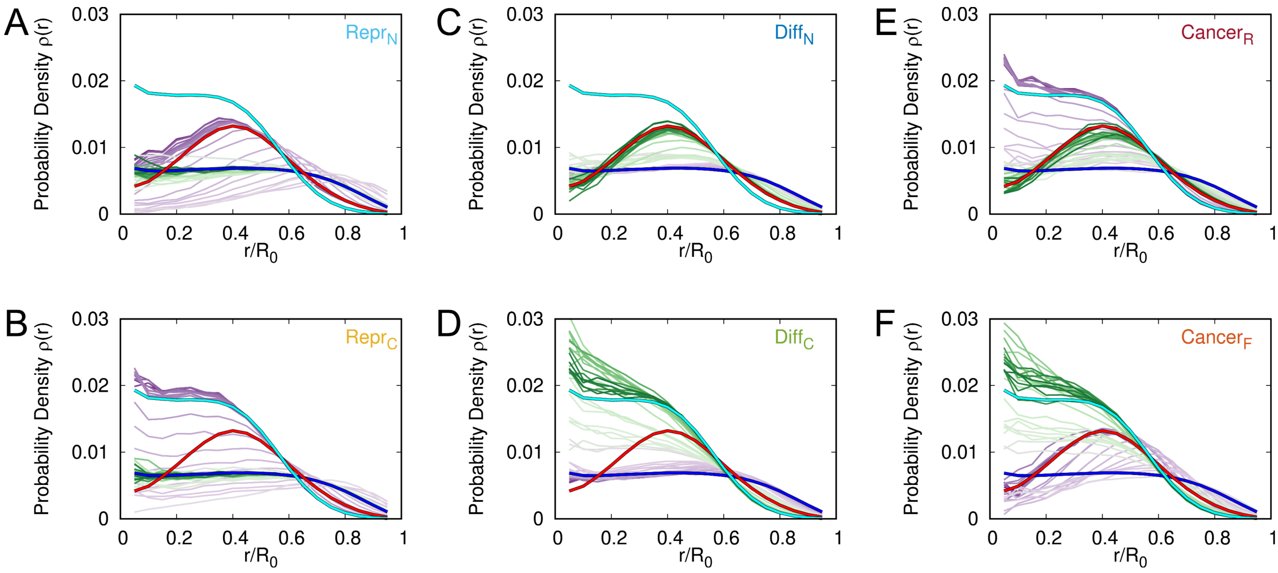


Figure S4: The change of the radial density $\rho(r)$ in the chromosome for the **loci in compartment A** during the processes of (A) normal cell reprogramming to ESC, (B) cancer cell reprogramming to ESC, (C) ESC differentiation to normal cell, (D) ESC differentiation to cancer cell, (E) cancer reversion to normal cell and (F) cancer formation from normal cell. Time is in the logarithmic scale.

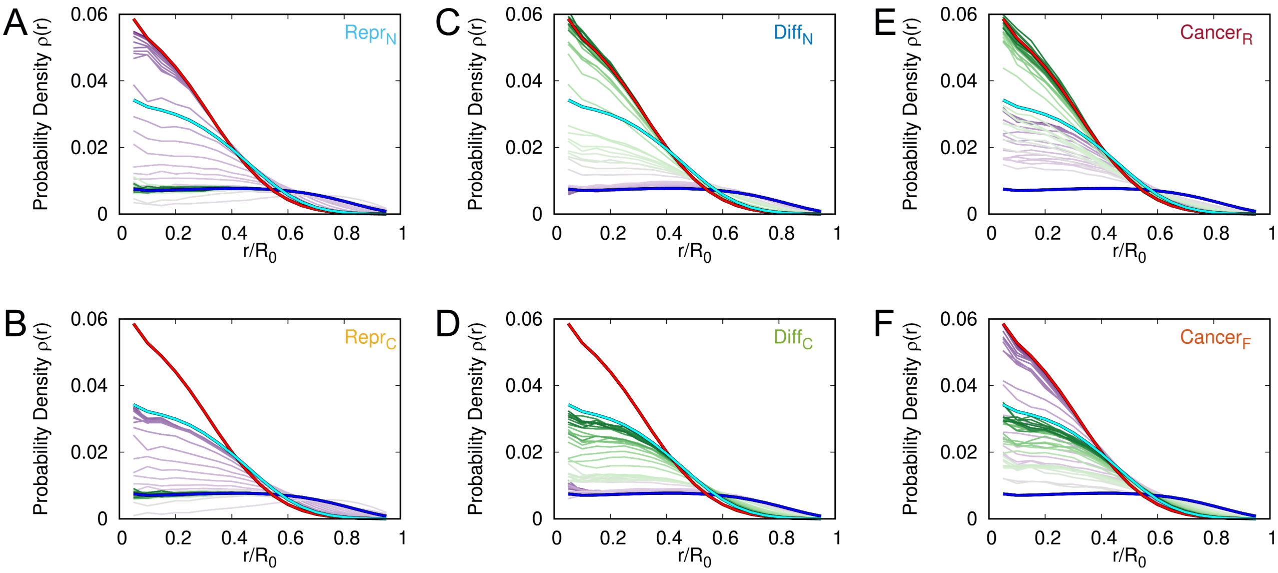


Figure S5: The change of the radial density $\rho(r)$ in the chromosome for the **loci in compartment B** during the processes of (A) normal cell reprogramming to ESC, (B) cancer cell reprogramming to ESC, (C) ESC differentiation to normal cell, (D) ESC differentiation to cancer cell, (E) cancer reversion to normal cell and (F) cancer formation from normal cell. Time is in the logarithmic scale.

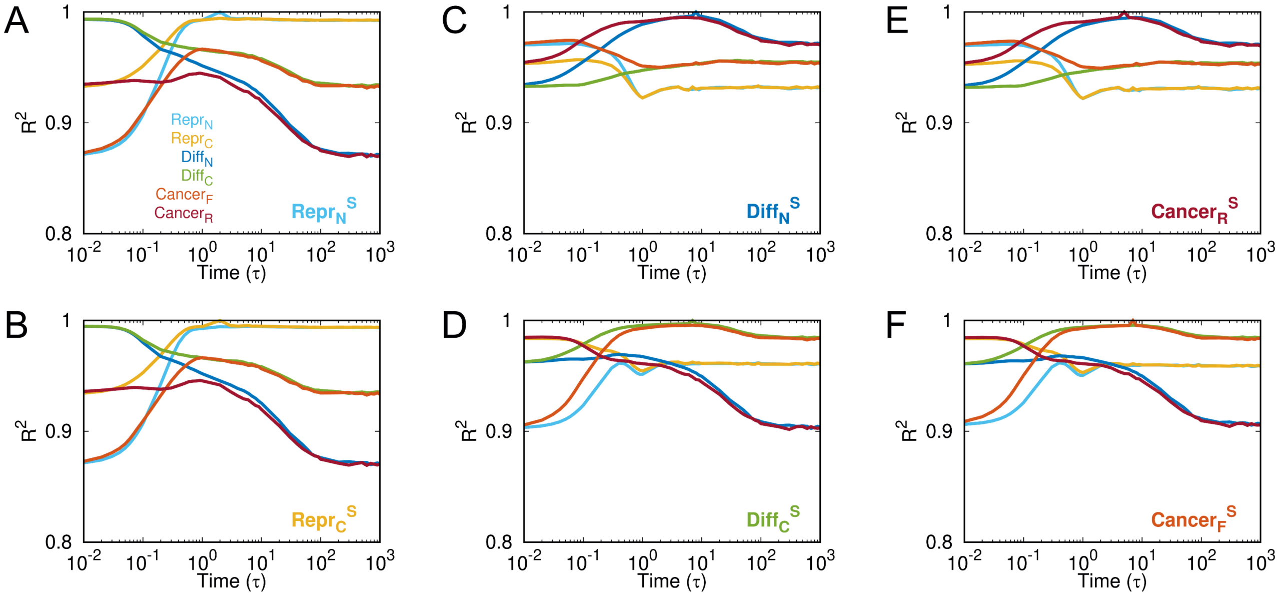


Figure S6: The coefficient of determination $R^2$ between contact probability maps in chromosome at the transient states and states during the processes of (A) normal cell reprogramming to ESC, (B) cancer cell reprogramming to ESC, (C) ESC differentiation to normal cell, (D) ESC differentiation to cancer cell, (E) cancer reversion to normal cell and (F) cancer formation from normal cell.

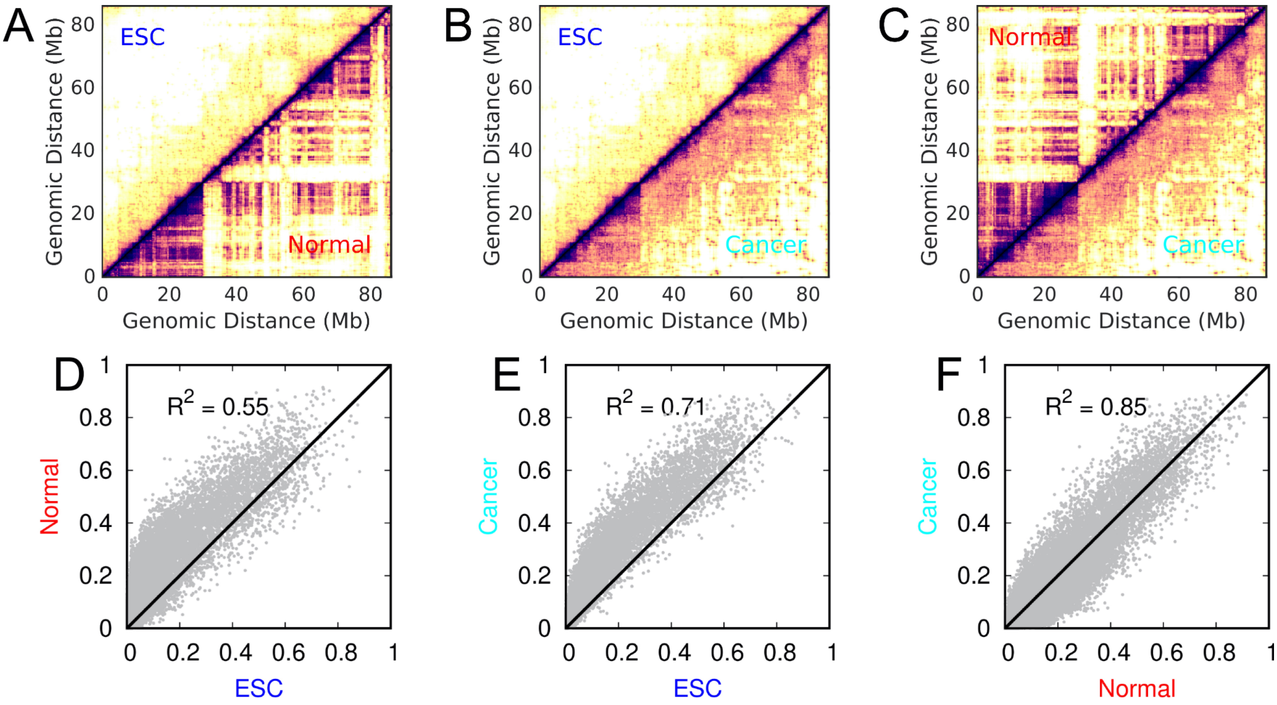


Figure S7: Comparisons of contact probability formed in chromosome among ESC, normal and cancer cells. The comparisons are shown in terms of (A-C) contact maps and (D-E) $R^2$ calculated between the contact probability of two cell states.

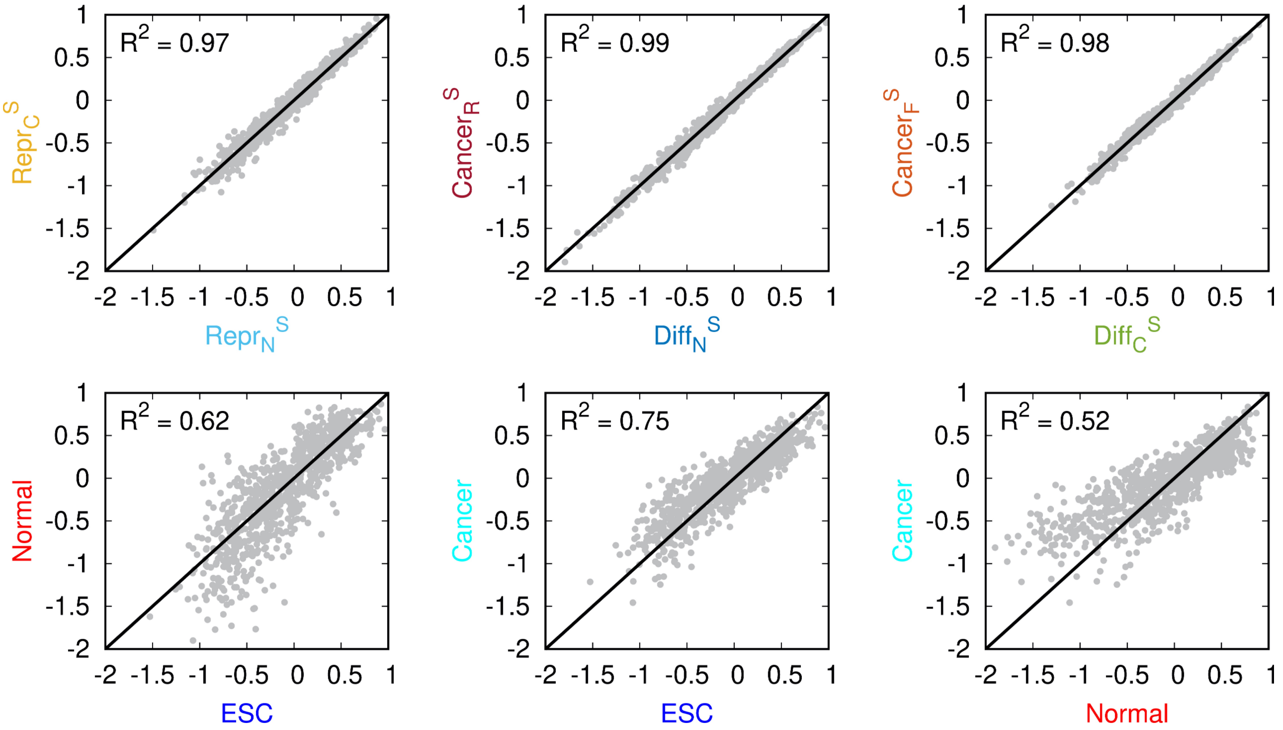


Figure S8: The coefficient of determination $R^2$ calculated between the insulation score profiles of two cell states. The insulation score signifies the TAD boundary formation [1].

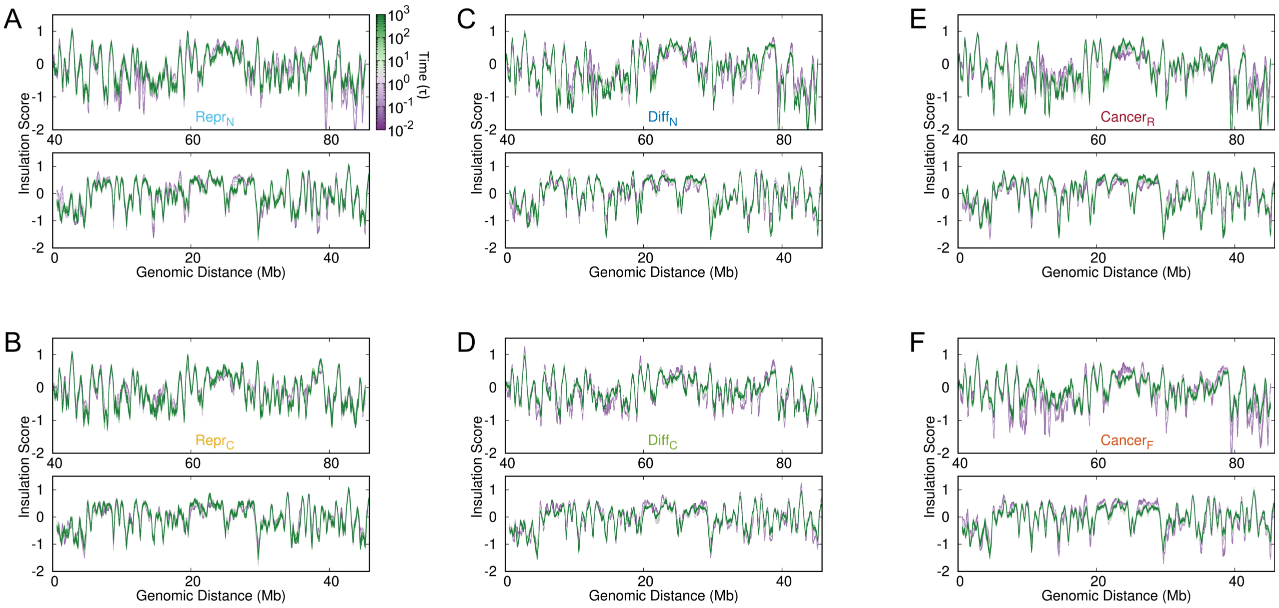


Figure S9: The change of insulation score of the chromosome during the processes of (A) normal cell reprogramming to ESC, (B) cancer cell reprogramming to ESC, (C) ESC differentiation to normal cell, (D) ESC differentiation to cancer cell, (E) cancer reversion to normal cell and (F) cancer formation from normal cell. Time is in the logarithmic scale.

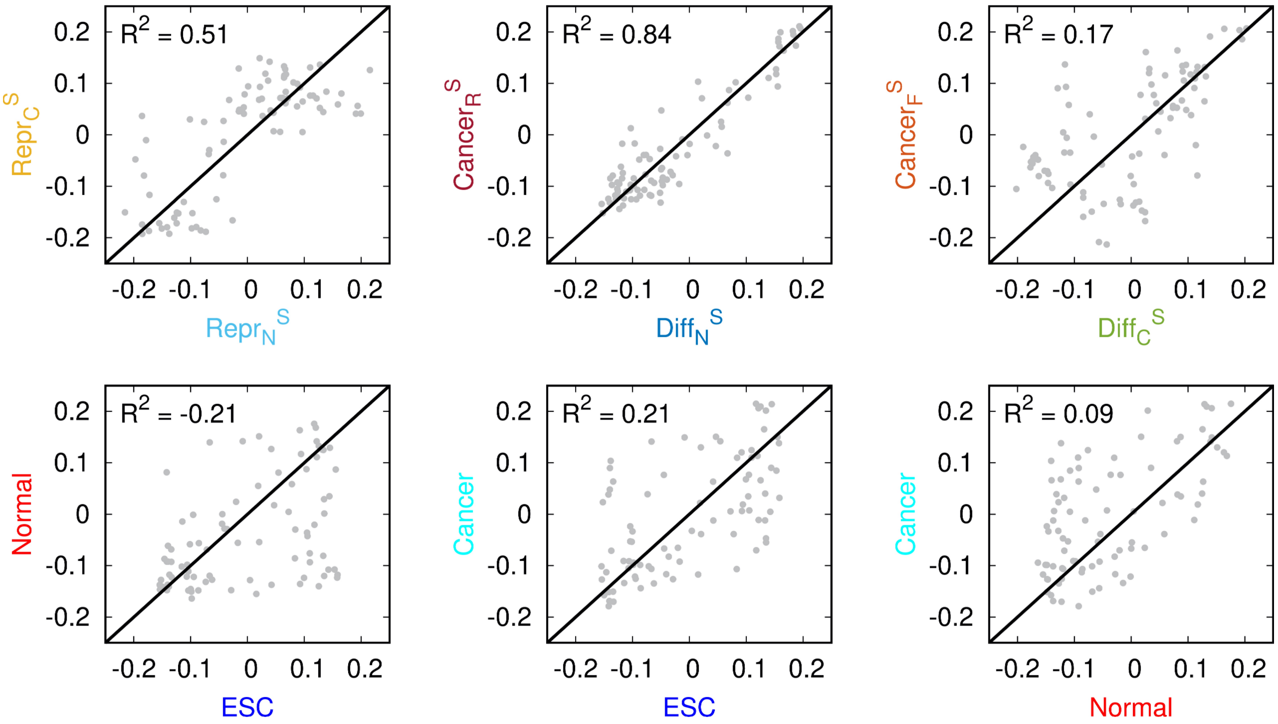


Figure S10: The coefficient of determination $R^2$ calculated between the compartment profiles of two cell states.
